# Truncated titin is integrated into the human dilated cardiomyopathic sarcomere

**DOI:** 10.1101/2023.02.08.527678

**Authors:** Dalma Kellermayer, Hedvig Tordai, Balázs Kiss, György Török, Dániel M. Péter, Alex Ali Sayour, Miklós Pólos, István Hartyánszky, Bálint Szilveszter, Siegfried Labeit, Ambrus Gángó, Gábor Bedics, Csaba Bödör, Tamás Radovits, Béla Merkely, Miklós S.Z. Kellermayer

## Abstract

Heterozygous (HET) truncating mutations in the TTN gene (TTNtv) encoding the giant titin protein are the most common genetic cause of dilated cardiomyopathy (DCM). However, the molecular mechanisms by which TTNtv mutations induce DCM are controversial. Here we investigated 127 clinically identified DCM human cardiac samples with next-generation sequencing (NGS), high-resolution gel electrophoresis, Western blot analysis and super-resolution microscopy in order to dissect the structural and functional consequences of TTNtv mutations. The occurrence of TTNtv was found to be 15% in the DCM cohort. Truncated titin proteins matching, by molecular weight, the gene-sequence predictions were detected in the majority of the TTNtv samples. The total amount of expressed titin, which includes the truncated fragments, was comparable in the TTNtv+ and TTNtv-samples, indicating that titin haploinsufficiency is not the leading cause of the molecular pathogenesis. Proteomic analysis of washed cardiac myofibrils and Stimulated Emission Depletion (STED) super-resolution microscopy of myocardial sarcomeres labeled with sequence-specific anti-titin antibodies revealed that truncated titin is structurally integrated in the sarcomere. Sarcomere length-dependent anti-titin epitope position, shape and intensity analysis pointed at structural defects in the I/A junction and the M-band of TTNtv+ sarcomeres, which may contribute, via faulty mechanosensor function, to the development of manifest DCM.

## Introduction

The giant myofilament titin is the third most abundant sarcomeric protein besides actin and myosin^1^. A single titin molecule spans half of the sarcomere from the Z-disk to the M-line^2^. Titin’s main function is to provide passive stiffness to striated muscle^3^, but it also plays a prominent role in sarcomere development and assembly^4^. Recent next-generation sequencing data have shown that mutations in the TTN gene, which encodes titin, are associated with skeletal and cardiac myopathies^5–7^. Among these, the most prevalent is dilated cardiomyopathy (DCM)^6^. DCM is characterized by ventricular and atrial enlargement and reduced ventricular systolic function^8^. The leading genetic cause of DCM is heterozygous (HET) truncating variant mutations in the TTN gene (TTNtv) which account for ~ 15-25% of the familial cases^6, 9, 10^. Most of the TTNtvs are nonsense and frameshift mutations, but splicing and copy-number mutations occur as well^6, 11^. TTNtvs are over-represented in the A-band section of titin^6, 9^ that consists of constitutively expressed exons. This section is functionally inextensible and acts as a molecular ruler for thick filament assembly^12^. Interestingly, TTNtv mutations also appear in ~1% of the healthy population, mainly in non-constitutive exons of the I-band, indicating a profuse splicing mechanism in titin’s I-band^9^.

The molecular mechanisms underlying TTNtv-induced DCM are still highly controversial. Multiple pathways have been proposed, including haploinsufficiency, poison peptide mechanism and perturbation of cardiac metabolism^10, 13, 14^. Moreover, it has been suggested that the TTNtv mutation itself is often insufficient to induce a phenotype, but an additional factor, such as a second gene mutation, pregnancy or an environmental stressor (e.g., alcohol, hypertonia, chemotherapy), is required to evoke clinical manifestation of the disease^15–21^. Recent studies by Fomin *et al*^22^ and McAfee *et al*^23^ demonstrated, for the first time, the presence of truncated titin proteins in human cardiac samples of dilated cardiomyopathy and suggested that titin haploinsufficiency may play a key role in the pathogenesis of DCM. Furthermore, Fomin *et al* showed that the truncated protein is not incorporated into the sarcomere but is accumulated in intracellular aggregates that act as toxic agents and impair protein quality control, supporting the poison peptide mechanism^22^. By contrast, McAfee *et al* suggested possible sarcomeric integration of the truncated protein^23^. Furthermore, experiments on human-induced pluripotent stem cell-derived cardiomyocytes (hiPSC-CMs) containing TTNtv showed impaired contractility relative to healthy controls^13, 24^. Overall, these findings suggest that the truncated protein is likely integrated into the sarcomere and further support the poison peptide mechanism. However, the sarcomeric presence and arrangement of truncated titin has not yet been detected directly in the myocardium of DCM patients.

Here, we investigated titin truncating variant mutations in a human cohort of DCM containing 127 patients. We analyzed the gene sequence of the samples to identify the location of titin truncation, and investigated the protein expression profiles to reveal the corresponding protein products. The sarcomeric arrangement of truncated titin was explored with super-resolution microscopy on myocardial samples labeled with sequence-specific antibodies and exposed to mechanical stretch. We found that truncated titin is structurally integrated in the sarcomere and causes small, albeit significant structural and functional disturbances which are the likely contributors to the pathway towards DCM.

## Methods

### Sample collection and handling

Human myocardial tissue samples were obtained from the Transplantation Biobank of the Heart and Vascular Center at Semmelweis University, Budapest, Hungary. The sample collection procedure was reviewed and approved by institutional and national ethics committees (permission numbers: ETT TUKEB 7891/2012/EKU (119/ PI/12.) and TUKEB 73/2005.). Informed consent was obtained from the patients prior to sample collection, in accordance with the Declaration of Helsinki. Myocardial septum samples were collected from 127 clinically identified, end-stage DCM patients undergoing orthotopic heart transplantation (HTx). The samples were surgically dissected from the explanted, diseased hearts of the recipients. The septum samples were immediately snap-frozen in liquid nitrogen under sterile conditions and stored at −80 °C for further measurements and analyses. Echocardiographic data recorded prior to surgery were obtained from our Transplantation Biobank database. The sample ID numbers shown in the images are the patients’s ID numbers from the Heart Transplantation registry.

### Gene sequencing

Genomic DNA was isolated from 25 mg of frozen septum samples by using the QIAamp DNA Mini Kit (Qiagen, Hilden, Germany) according to the manufacturer’s recommendation. Nextgeneration sequencing of the purified DNA (50 ng) was performed by using the Illumina TruSight Cardio library preparation kit^25^, with the libraries sequenced on a MiSeq instrument (Illumina Inc., San Diego, CA, USA). Quality control of the raw *fastq* files was performed with the FastQC (v0.11.9) and MultiQC (v1.9) algorithms^26, 27^. Further bioinformatic analyses were carried out with the Genome Analysis Toolkit (GATK) Best Practices of Germline Short Variant Discovery of the Broad Institute^28^. Sequence alignment to the hg19 genome version was performed by using the Burrows-Wheeler Aligner (BWA v.0.7.17), then the mapped reads underwent duplicate-marking (GATK-MarkDuplicatesSpark tool) and Base Quality Score Recalibration (GATK-BQSR tool)^29^. The variant calling step was performed for every sample (GATK-Haplotype Caller). The GVCF files produced for every analyzed sample were consolidated into a single file (GATK-GenomicsDBImport) in order to perform joint genotyping (GATK-GenotypeGVCF). This cohort-wide view facilitated highly sensitive detection of variants even at difficult genomic sites. Final filtering was performed with hardfiltering (with the following parameters: MIN_QD= 2MAX_FS=60). The filtered variants were annotated with dbSNP (v. human_9606_b151), COSMIC (v92), Ensembl-Variant Effect Predictor and ClinVar (v 2020.07.17) databases.

### Protein solubilization

10-15 mg pieces of the myocardial septum samples were homogenized in glass Kontes Dounce tissue grinders under liquid nitrogen. After 20 min of incubation at −20°C, the samples were solubilized at 60 °C for 15 min in 50% urea buffer (8 M Urea, 2 M Thiourea, 50 mM Tris-HCl, 75 mM DTT, 3% SDS and 0.03% Bromophenol blue, pH 6.8) and 50% glycerol containing protease inhibitors (0.04 mM E64, 0.16 mM Leupeptin and 0.2 mM PMSF). All solubilized samples were centrifuged at 13,000 rpm for 5 min, aliquoted, flash frozen in liquid nitrogen and stored at −80°C^30^.

### Titin isoform analysis

Titin expression level was determined by using 1% sodium-dodecyl-sulfate (SDS) – agarose gel electrophoresis^31^, performed at 16 mA/gel for 3.5 h. Subsequently, the gels were stained overnight with SYPRO Ruby Protein Gel Stain (Thermo Fischer Scientific, Waltham, MA, USA), then digitized with a Typhoon-laser scanner (Amersham BioSciences, Little Chalfont, Buckinghamshire, United Kingdom). ImageJ (National Institutes of Health, Bethesda, MD, USA) was used to analyze the optical density of the titin bands. Relative titin isoform ratio (N2BA/N2B ratio) was calculated from the integrated band densities. Relative content of full-length titin (T1) that includes N2BA and N2B was normalized to the myosin heavy chain (MyHC). T2 (titin’s proteolytic degradation product) was normalized to T1. Truncated proteins detected on the gels were normalized to T1.

### Detection of truncated titin products

To determine whether the suspected band visible on the gel is indeed a truncated titin product, we performed Western blot analysis. The samples were separated on a 0.8% SDS-agarose gel, then transferred onto PVDF membrane (Hybond-LFP, Amersham BioSciences, Little Chalfont, Buckinghamshire, United Kingdom) by using a semi-dry blotter (Trans-Blot Cell, Bio-Rad, Hercules, CA, USA). To differentiate the truncated product from T2, we used two antibodies, each detecting the terminal regions of titin. The blots were probed with anti-T12 (binds near titin’s N-terminus, kindly provided by Dieter O. Fürst, University of Bonn, Germany, dilution 1:1000)^32^ and anti-M8M10 (binds near titin’s C-terminus, Myomedix Ltd., Mannheim, Germany, dilution 1:1000) primary antibodies overnight at 4°C, followed by secondary CyDye-conjugated antibodies (Amersham BioSciences, Little Chalfont, Buckinghamshire, United Kingdom). Subsequently, the blots were digitized with a Typhoon laser-scanner. Relative expressions of the proteins were analyzed by using ImageJ.

### Preparation of myofibril suspension

Myofibril suspension was prepared based as previously described^33^. Briefly, 2 mL of ice-cold permeabilization solution (10 mM Tris (pH 7.1), 132 mM NaCl, 5 mM KCl, 1 mM MgCl2, 5 mM EGTA, 5 mM DTT, 10 mM NaN3, 20 mM 2,3-butanedione-monoxime (BDM), 1% Triton X-100) containing protease inhibitors (0.04 mM E64, 0.16 mM Leupeptin and 0.2 mM PMSF) was added to previously frozen, small pieces of septum samples (total weight 15 mg). Then the samples were incubated on a 360° rotating shaker for 3h at 4°C. Subsequently, the permeabilized samples were rinsed in washing solution (same as permeabilization solution, but without Triton X-100 and BDM) for 15 min. To prepare a myofibril suspension, the samples were transferred into 1 mL ice-cold fresh washing solution and homogenized with a MT-30K Handheld Homogenizer for 10-15 sec at 27,000 rpm (Hangzhou Miu Instruments Co. Ltd, Hangzhou, Zhejiang, China). The myofibrils were pelleted by applying low-speed centrifugation followed by resuspension and washing. The centrifuge and washing steps were performed at least twice. Subsequently, the pellet was solubilized in 50% urea buffer and 50% glycerol with inhibitors (1:1) and was incubated for 15 min at 60 °C. Additional 1:3 and 1:10 dilutions of the myofibril suspension:urea buffer solution were prepared. The solubilized myofibril samples were separated on 1% SDS-agarose gel.

### Super-resolution microscopy

Pieces of flash-frozen DCM ^TTNtv−^ (n=3) and DCM^TTNtv+^ (n=3) left ventricular cardiac muscle were dissected in relaxing solution (40 mM BES, 10 mM EGTA, 6.56 mM MgCl2, 5.88 mM Na-ATP, 1 mM DTT, 46.35 mM K-propionate, 15 mM creatine phosphate, pH 7.0) containing 1% (w/v) Triton X-100 and protease inhibitors (0.1 mM E64, 0.47 mM leupeptin and 0.25 mM PMSF). The dissected cardiac muscle pieces were skinned overnight at 4 °C in relaxing solution containing 1% Triton X-100 and protease inhibitors, washed thoroughly for at least 5 hours with relaxing solution and used immediately for experiments. Myofibril bundles were prepared and stretched from the slack length to different degrees (~ 40% to 70%), then fixed with 4% (v/v) formaldehyde diluted with phosphate buffer at neutral pH. Fixed bundles were embedded in optimal cutting temperature (O.C.T) compound and frozen immediately in 2-methylbutane pre-cooled by liquid nitrogen. 8-μm-thick cryosections were then cut with a Microm Cryo Star HM 560 cryostat (Thermo Fisher Scientific, Waltham, MA, USA) and mounted on microscope slides (Superfrost UltraPlus, Thermo Fisher Scientific). Tissue sections were permeabilized in 0.2% Triton X-100/PBS for 20 minutes at room temperature, blocked with 2% BSA and 1% normal donkey serum in PBS for 1 hour at 4 °C, and incubated overnight at 4 °C with primary antibodies and phalloidin diluted in blocking solution. The primary antibodies included: a rabbit polyclonal anti-titin MIR (TTN-7; www.myomedix.com, 0.4 μg/mL, 1:300 dilution), a rabbit polyclonal anti-titin A170 (TTN-8; www.myomedix.com, 0.7 μg/mL, 1:250 dilution) and AlexaFluor-488-conjugated phalloidin (Invitrogen, Waltham, MA, USA, A-12379, 66 μmol/L, 1:500 dilution). Sections were then washed with PBS for 2×30 minutes and incubated with Abberior STAR580 goat anti-rabbit IgG (Abberior GmbH, Göttingen, Germany) (1:250 dilution) and AlexaFluor-488-conjugated phalloidin (Invitrogen A-12379, 66 μmol/L, 1:500 dilution). The sections were then washed with PBS for 2×15 minutes and covered with number 1.5H high precision coverslips (Marienfeld Superior, Lauda-Königshofen, Germany) by using ProLong Diamond (Thermo Fisher Scientific) for 24 hours to harden. Stimulated Emission Depletion (STED) microscopy was performed by using an Abberior Expert Line microscope (Abberior GmbH). For the excitation of conjugated phalloidin and titin epitopes, 488 nm and 560 nm laser illumination sources were used, respectively. For STED imaging of the different titin epitopes labeled with STAR580 dye, a 775 nm depletion laser was utilized. Images were acquired by using a Nikon CFI PL APO 100x (NA=1.45) oil immersion objective coupled with avalanche photodiode detectors with spectral detection capabilities. Deconvolution of the recorded STED images was performed by using Huygens Professional software (SVI) using the theoretical point spread function (PSF) of the imaging objective. Fluorescence intensity plot profiles were generated with Fiji (based on ImageJ v. 1.52, NIH, USA). Plot profiles were fitted with Gaussian curves to determine the epitope peak position, height and full width at half maximum (FWHM) using Fityk 1.3.0 software. A-band width was determined from the MIR epitope positions across the Z-disk. Fluorescence intensity normalization of signals of the A170 titin epitope was performed on carefully pre-selected plot profiles. For this purpose, background intensity-corrected plot profiles were collected with averaging across a thick line to compensate for labeling inhomogeneity along the epitope lines. Plot profiles were discarded in case the intensity fluctuation of either the MIR or the A170 epitopes across the M-line exceeded 20% of the average intensity of the respective epitope. Fluorescence intensity was calculated from the peak height of the fitted Gaussians.

### Statistics

Statistical analysis was performed by using GraphPad Prism 8 (GraphPad Software, Inc.). Continuous-variable statistical data are shown as mean ± standard error of the mean unless stated otherwise. Differences between groups were considered to be statistically significant at a probability value of p<0.05. Two-tailed t-test was used when comparing the means of two groups, and Welch’s correction was applied in the case of unequal variances between the two groups. Normality of statistical distribution was checked with the Shapiro-Wilk test. Mann-Whitney U-test was used in statistical comparison of data for which normal distribution could not be assumed. In order to increase the statistical power of the tests, equal or close to equal sample size was applied within independent groups. Linear regression analysis was used to fit and compare sarcomere-length-dependent epitope distance and fluorescence intensity data obtained from super-resolution microscopy. Sample randomization as well as blinding of the investigator was applied in the case of the analysis of the super-resolution microscopy images. For detailed statistical information, see **Supplementary Tables**.

## Results

### Patient data and gene sequencing: Identification of a TTN-DCM cohort

We screened 127 myocardial samples from explanted hearts of clinically identified DCM patients (**Table 1** and **Supplementary Table 1**) for potentially pathogenic genetic variants by using targeted exome sequencing (next-generation sequencing, NGS). The DNA libraries were prepared so as to identify variants in 174 genes associated with inherited cardiac conditions^25^. 50 out of the 174 genes are associated with DCM, whereas the remaining cases are implicated in inherited arrhythmias, other cardiomyopathies, aortopathies and familial hypercholesterolemia, respectively. We identified 35635 variants by using the GATK pipeline^28^, from which 13815 were found in DCM-associated genes and 4428 in the titin (TTN) gene (**Supplementary Table 2**). The variants were annotated by comparing with the dbSNP, COSMIC and ClinVar databases. Based on the annotations and newly identified frameshift and nonsense variants, we found potentially pathogenic heterozygous mutations in 44 samples (**Supplementary Table 2**). In 35 samples (27.5%) the mutations were in DCM genes. We identified 19 TTNtv (titin truncating variant, 15%), 4 LMNA (lamin A), 4 DSP (desmoplakin), 2 BAG3, and one each of FKTN, LAMA2, MYBPC3, MYH6, MYH7, PLN, RBM20 and TNNI3 variants (**Supplementary Tables 2-3**). Out of the 19 pathogenic, likely pathogenic and new heterozygous TTNtv mutations eight were frameshift and eleven were nonsense mutations (**Supplementary Table 3)**. All variants were located in the constitutively expressed I/A junction, A- and M-band regions of titin (**Figure 1, Supplementary Table 4**). In two TTNtv samples we identified additional potentially pathogenic mutations in the RAF1, and TRPM4 genes (**Supplementary Table 3**). Based on the NGS data we divided our samples into DCM samples with (DCM^TTNtv+^, n=19) or without (DCM^TTNtv−^, n=108) titin truncation. We evaluated the echocardiographic data of the patients recorded prior to the transplantation to examine any TTNtv-associated phenotype (**Table 1**). Although more men had TTNtv mutations similarly to the finding of others^6^, we did not reveal any genotype-associated phenotype severity in our DCM population. None of the functional parameters showed differences between the two groups.

**Table 1.** Echocardiographic data of DCM patients with (DCM^TTNtv+^) and without (DCM^TTNtv−^) titin truncation variant. Data are shown as mean±SD. LV: left ventricle; RV: right ventricle; EDD: end-diastolic diameter; IVSDd: intraventricular septum diameter (diastole); LA-LD: left atrial longitudinal diameter; LA-HD: left atrial horizontal diameter; RA-LD: right atrial longitudinal diameter; RA-HD: right atrial horizontal diameter; AAD: ascending aorta diameter; EF: ejection fraction (Simpson method); TAPSE: tricuspid annular plane systolic excursion. Note that the variation in sample size is due to the heterogeneity in end-stage clinical profiling.

| Patient and echocardiographic data | DCM<sup>TTNtv-</sup> | DCM<sup>TTNtv+</sup> | p-value |
| --- | --- | --- | --- |
| Age at HTX (years) | 50±10 (n=108) | 49±11 (n=19) | 0.83 |
| Sex (F,%) | 26/82 (24%) (n=108) | 1/18 (5.3%) (n=19) |  |
| LV-EDD (mm) | 70.8±11.30 (n=95) | 70.0±10.6 (n=17) | 0.78 |
| RV-EDD (mm) | 41.9±7.75 (n=78) | 39.0±5.7 (n=14) | 0.11 |
| IVSDd (mm) | 10.2±7.4 (n=77) | 9.1±1.6 (n=13) | 0.23 |
| LA-LD (mm) | 56±11 (n=88) | 56.0±10.7 (n=16) | 0.99 |
| LA-HD (mm) | 56.1±10.7 (n=72) | 58.0±11.5 (n=12) | 0.61 |
| RA-LD (mm) | 49.9±15.7 (n=87) | 50.1±6.9 (n=16) | 0.93 |
| RA-HD (mm) | 54.6±10.5 (n=67) | 55.3±9.8 (n=11) | 0.85 |
| AAD (mm) | 31.3±7.9 (n=72) | 30.4±4.2 (n=13) | 0.52 |
| EF (%) | 22.3±6.5 (n=97) | 21.2±5.6 (n=15) | 0.49 |
| TAPSE (mm) | 15.7±4.6 (n=88) | 16.1±5.1 (n=14) | 0.76 |

**Figure 1.**
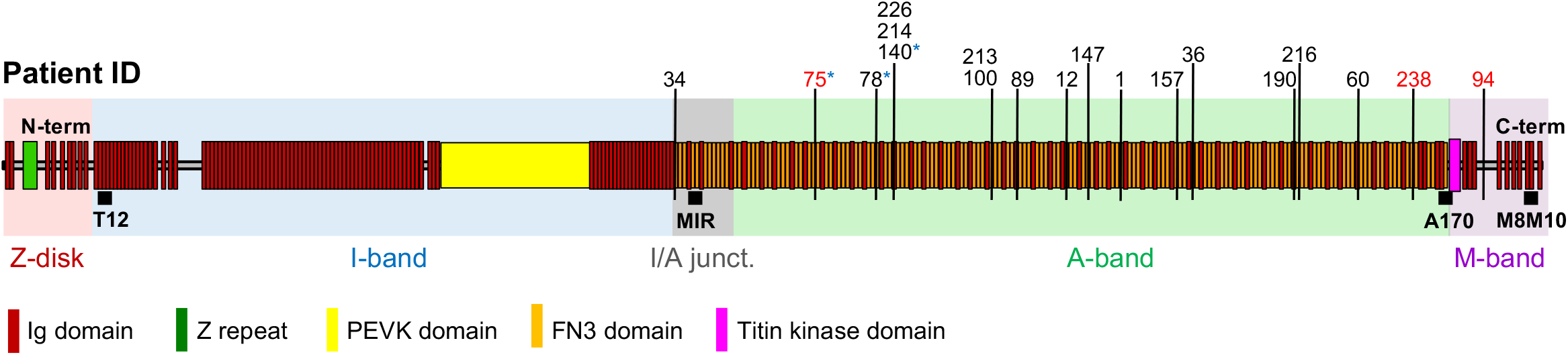
Titin domain structure and location of truncating variants. Layout of the human titin isoform IC (NCBI Reference Sequence NP_001254479) comprising of 35991 amino acids. Next-generation sequencing (NGS) revealed 18 DCM samples with TTNtv, labeled according to the registry patient number. The truncating variants were over-represented in the A-band region of titin. Patient #34 had a truncating variant in the I/A junction, patient #94 had the TTNtv in the M-band region. Red and starred numbers indicate samples used for myofibril protein composition and structural (STED microscopy) analyses, respectively. Black squares indicate the binding location of monoclonal antibodies used in STED microscopy.

### Protein analysis detects TTNv subspecies and perturbed stoichiometries

Next, we analyzed the titin expression profiles of the myocardial samples with high-resolution gel electrophoresis. The N2BA/N2B titin isoform ratio was elevated in both the DCM^TTNtv−^ and DCM^TTNtv+^ groups compared to physiological ratios obtained from literature data^34^ (**Figure 2**). The full-length titin to myosin heavy chain ratio (T1/MyHC) was significantly decreased in the DCM^TTNtv+^ group. The ratio of the T2 fragment (a calpain-dependent proteolytic fragment that encompasses titin’s A-band section and a 100-200 kDa portion of its distal I-band section^35^) to full-length titin (T2/T1) was significantly increased in the DCM^TTNtv+^ samples (**Figure 2D**), suggesting that proteolytic activity may be increased in the DCM^TTNtv+^ myocardial tissue. However, the (T1+T2)/MyHC ratio did not differ significantly between the two groups (**Figure 2F**). We detected additional protein bands in the DCM^TTNtv+^ group (**Figures 2A and S1**), albeit not in all TTNtv+ samples. These proteins were observed on the gels at the most probable molecular weights calculated for the respective truncated titins from gene-sequencing data (**Supplementary Table 4**). Accordingly, we identify these bands as the protein products of the truncated titin genes. The relative expression of the truncated proteins to full-length titin (T1) was 0.19. Notably, upon adding the truncated protein quantity to the respective T1 (**Figure 2E**), there was no significant difference between the DCM^TTNtv+^ and DCM^TTNtv−^ samples. Furthermore, the integrated titin quantities (T1 + T2 + truncated titin) normalized to MyHC were essentially identical in DCM^TTNtv+^ and DCM^TTNtv−^.

**Figure 2.**
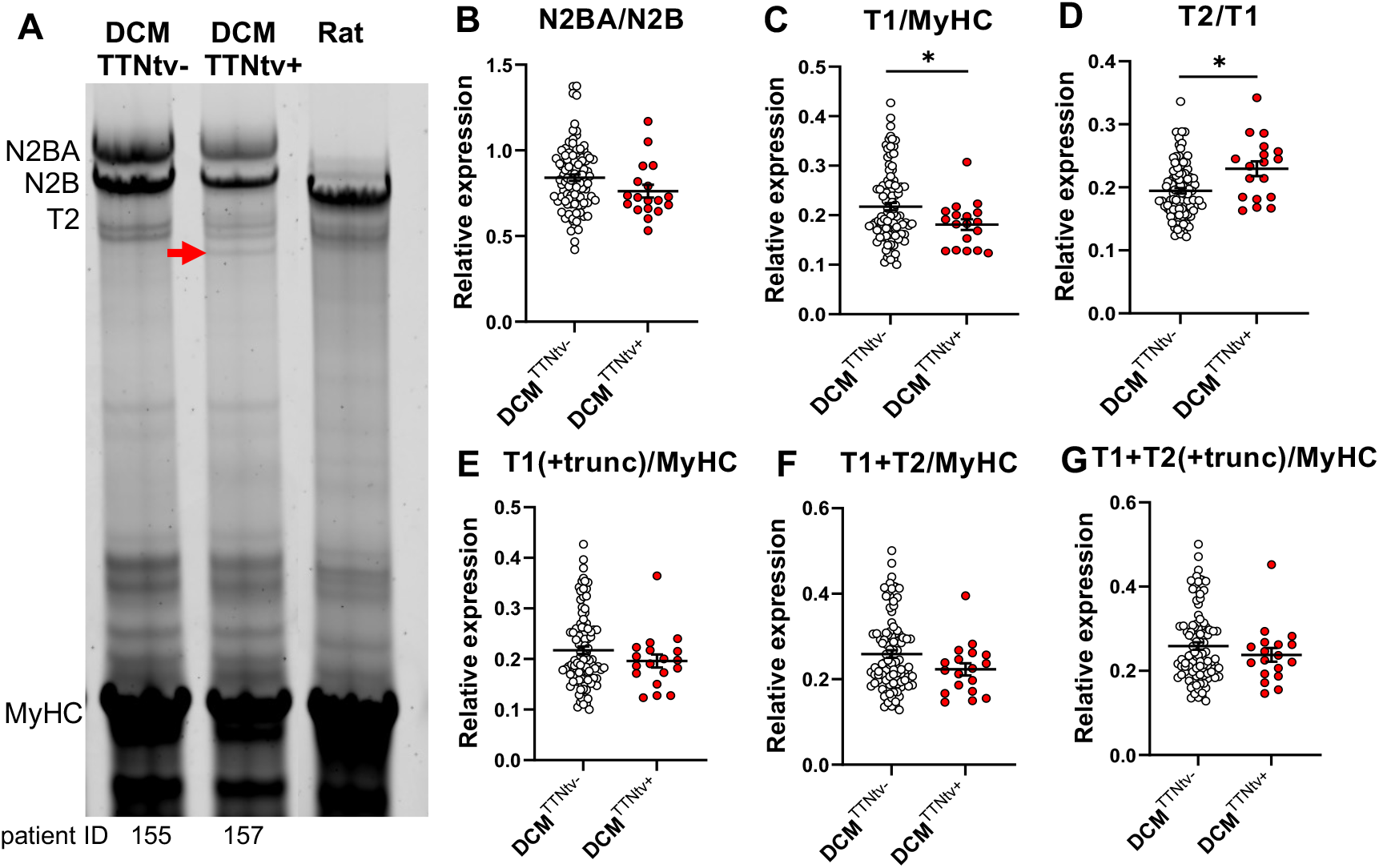
Titin isoform analysis. **A,** Representative image of gel electrophoresis demonstrating titin N2BA and N2B isoforms, titin’s proteolytic degradation product (T2), myosin heavy chain (MyHC) in TTNtv-, TTNtv+ human myocardium and rat myocardium. The red arrow indicates the truncated protein. Linear contrast adjustment also applied to the original image for better visualization. **B,** The N2BA/N2B ratio did not differ between TTNtv+ (n=18) vs. TTNtv-(n=98) DCM groups. **C,** Total titin (T1=N2BA+N2B) normalized to MyHC was significantly reduced in the DCM^TTNtv+^ samples. **D,** T2/T1 increased significantly in the TTNtv+ DCM samples. **E,F,G,** Although T1/MyHC was reduced in the TTNtv+ DCM group, no differences were seen in the samples if the truncated protein and T2 were added to T1. Values are mean±SEM, *p<0.05.

To experimentally test whether the additional protein bands are indeed truncated titins rather than the product of proteolysis (such as T2), we carried out Western-blot analysis by using sequence-specific antibodies targeting the C- and N-terminal regions of titin. The T12 antibody, which binds towards titin’s N-terminus, labeled the additional protein bands but not T2 (**Figure 3A**). By contrast, the M8M10 antibody, which binds near titin’s C-terminus, labeled T2, but not the additional protein bands (**Figure 3B**). Thus, we could differentiate the additional protein bands from T2, demonstrating that they contain titin’s N-terminal region, and proving that they indeed correspond to the protein products of the truncated titin genes.

**Figure 3.**
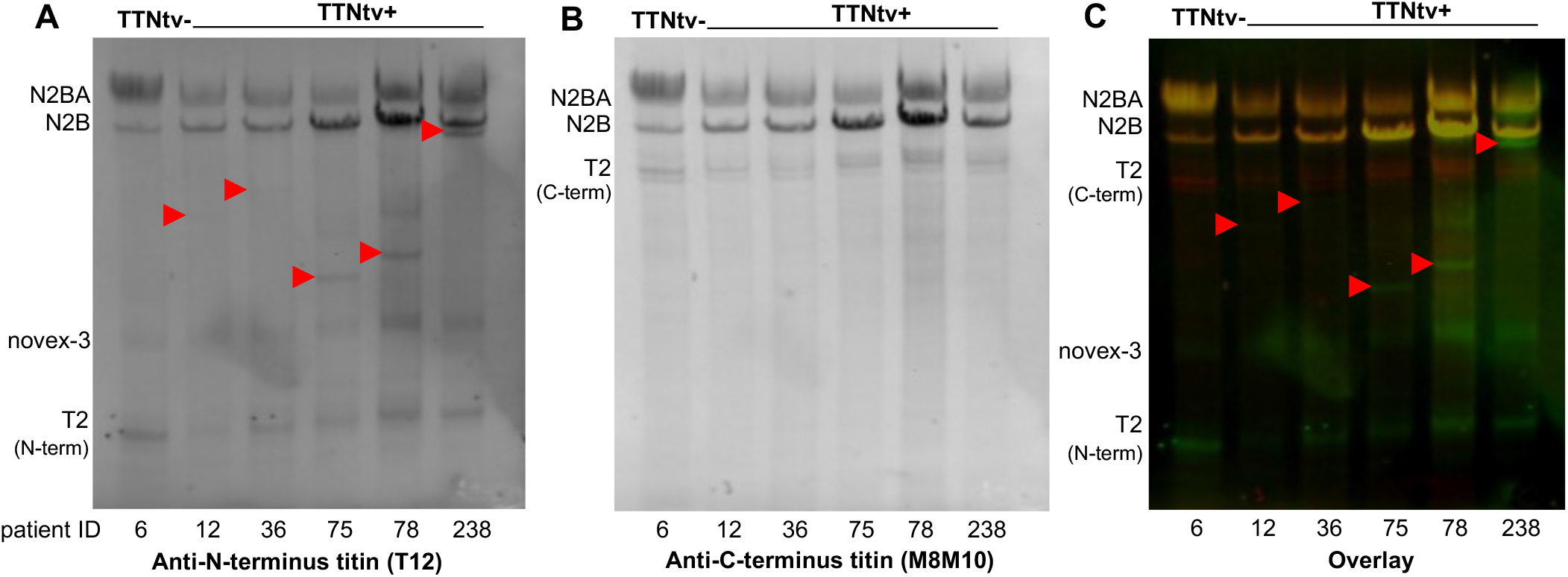
Differentiation of truncated titin from T2 by Western blot analysis. **A,** Anti-N-terminus antibody (T2) detected truncated titin, novex-3 and the N-terminus part of T2. **B,** No signal was detected at the molecular level of the truncated proteins with the anti-C-terminus titin antibody M8M10, but it labeled the C-terminus part of T2. **C,** The overlay image clearly differentiated the truncated titin from T2. Green: T12, red: M8M10.

To investigate whether the truncated titins are incorporated in the sarcomere rather than present in the bulk of the sarcoplasm, we carried out protein analysis of washed DCM^TTNtv+^ myofibrils (**Figures 4 and S2**). We were able to detect protein bands corresponding to the respective truncated titins in the gel electrophoretograms of washed DCM^TTNtv+^ myofibrils. Thus, we conclude that the truncated titin protein is incorporated in the cardiac muscle sarcomere.

**Figure 4.**
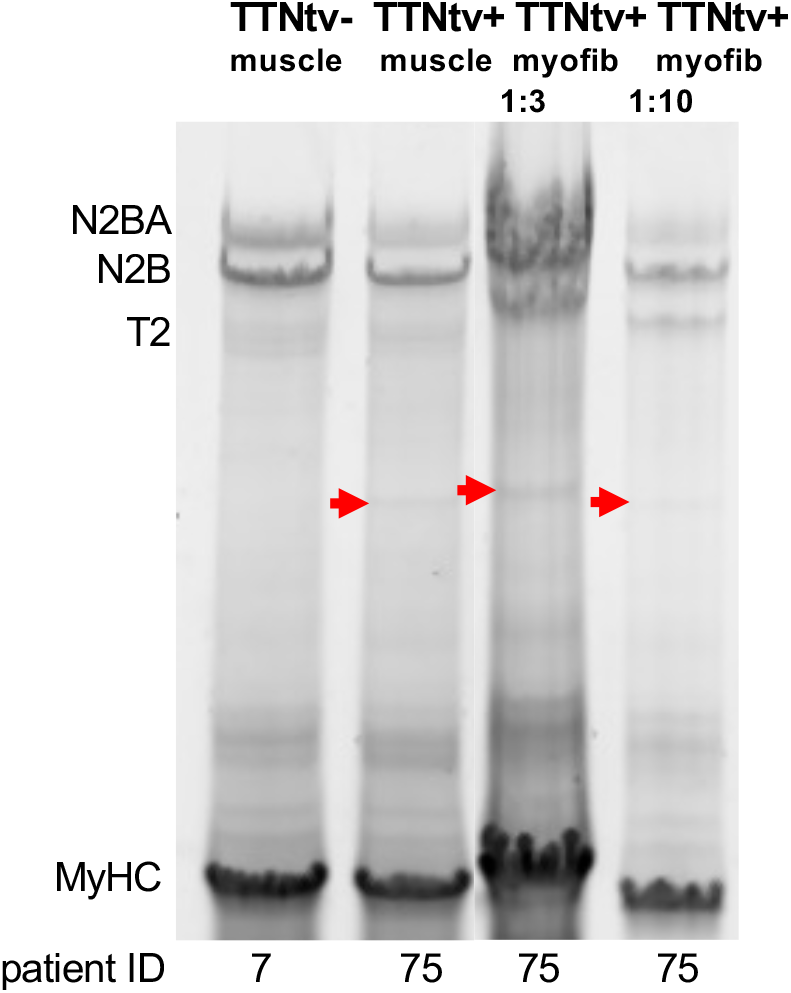
Skinned myofibrils contain the truncated titin. SDS-PAGE profile of washed myofibril pellets solubilized in urea buffer with 1:3 and 1:10 ratio. In contrast to the TTNtv-sample (patient #7), the TTNtv+ myofibril sample (patient #75) contains the truncated titin, indicating that the truncated protein is part of the sarcomere (see also **Figure S2**).

### Super-resolution microscopy detects altered A/I-junction and M-line widths

To explore whether and how truncated titin is structurally integrated in the sarcomere, we analyzed cardiac muscle samples, labeled with sequence-specific anti-titin antibodies (MIR and A170), by using super-resolution STED microscopy (**Figures 5–7**). We were unable to discern gross structural changes in the DCM^TTNtv+^ sarcomeres with respect to the DCM^TTNtv−^ samples (**Figure 5**). The A170 doublet (separated by ~140 nm) could be resolved in both groups with STED, but not with confocal microscopy (**Figure 6B bottom**). The relative intensity of the A170 epitope, normalized to the intensity of the MIR epitope of the same sarcomere, was significantly decreased in DCM^TTNtv+^ muscles (0.1765) compared to that in DCM^TTNtv−^ (0.2646), which is consistent with the missing epitope in truncated titin (**Figures 6A and B**). The ratio of these intensities is 0.667, which is in good agreement with the proteomic ratio of the expressed truncated and full-length titins in the respective DCM^TTNtv+^ sample (**Figure 7C**). Neither epitope doubling (**Figures 6A and B**), nor significant intensity differences (data not shown) were found in the case of the MIR epitope which is present in both the full-length and truncated titin.

**Figure 5.**
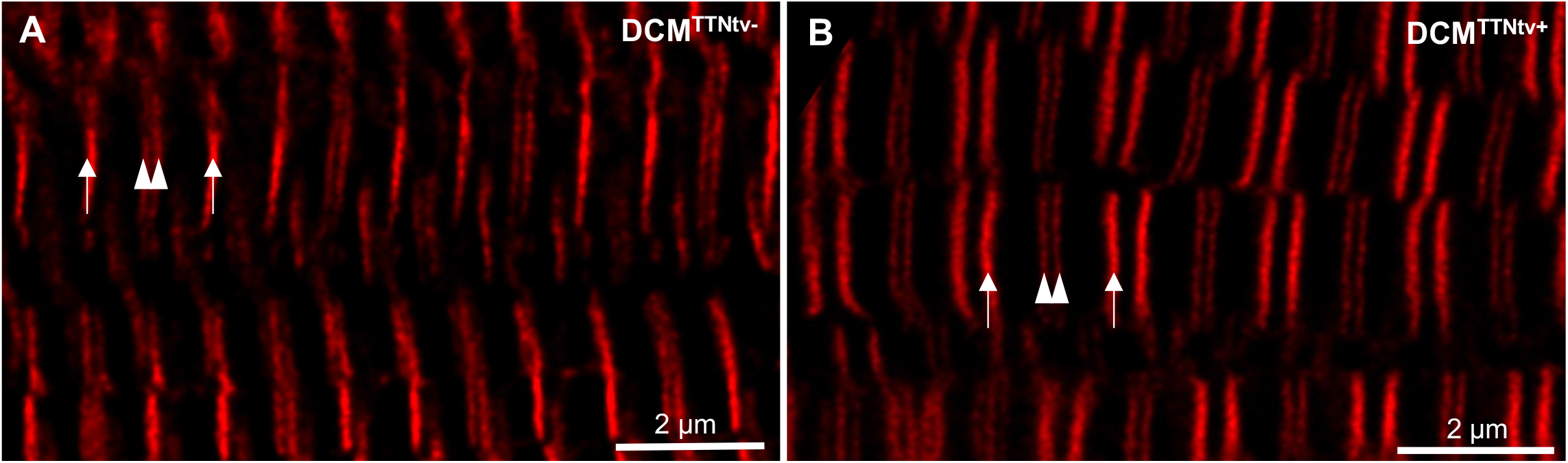
STED super-resolution microscopy of human DCM cardiac muscle samples. **A** and **B,** Representative STED images of DCM^TTNtv−^ and DCM^TTNtv+^ cardiac samples labeled with MIR and A170 antibodies, respectively. Arrows point at the MIR epitopes within one cardiac sarcomere, and arrowheads indicate the A170 epitopes near the M-line. Note that no gross structural alterations can be discerned in the DCM^TTNtv+^ sarcomeres.

**Figure 6.**
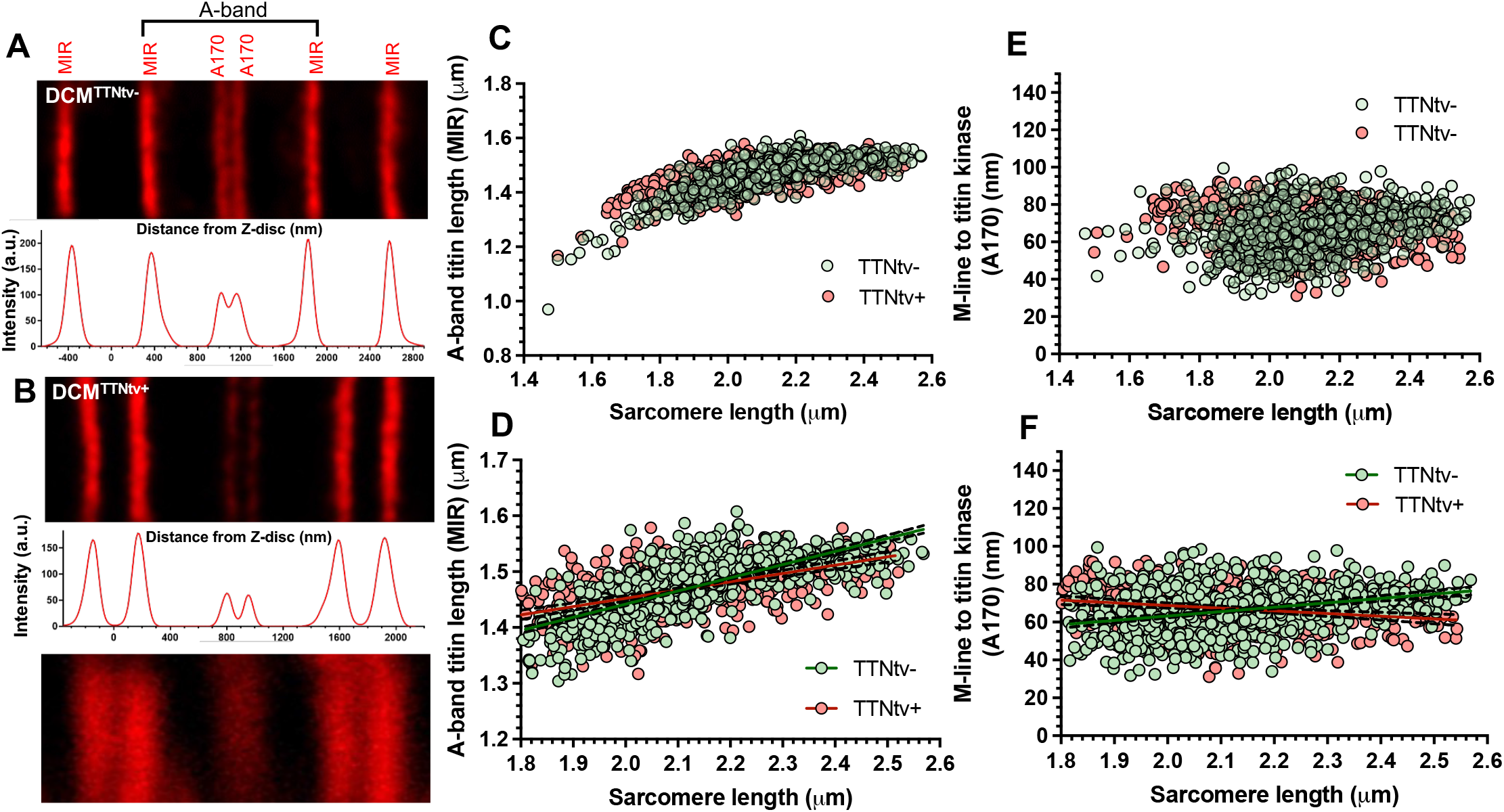
Titin epitope position and distance analysis. **A** and **B,** Representative STED images and plot profiles of DCM^TTNtv−^ and DCM^TTNtv+^ cardiac samples labeled with MIR and A170 anti-titin antibodies. Note the decreased A170 intensity in DCM^TTNtv+^ sarcomeres relative to that of the MIR epitope. The bottom panel in B is the respective confocal microscopic image, shown here to indicate the critical power of STED in resolving the separate A170 epitopes. **C**, A-band titin length (measured as the distance between two consecutive MIR epitopes bounding an A170 epitope doublet) increases in both DCM^TTNtv−^ and DCM^TTNtv+^ cardiac fibers when the sarcomeres are passively stretched. **D**, Regression analysis of the A-band titin length in the 1.8-2.6 μm sarcomere length range. **E**, M-line to titin kinase distance, calculated as the half value of the distance of two consecutive A170 epitopes, as a function of sarcomere length. **F**, Regression analysis of the sarcomere-length-dependent M-line to titin kinase distance in the 1.8-2.6 μm sarcomere length range.

**Figure 7.**
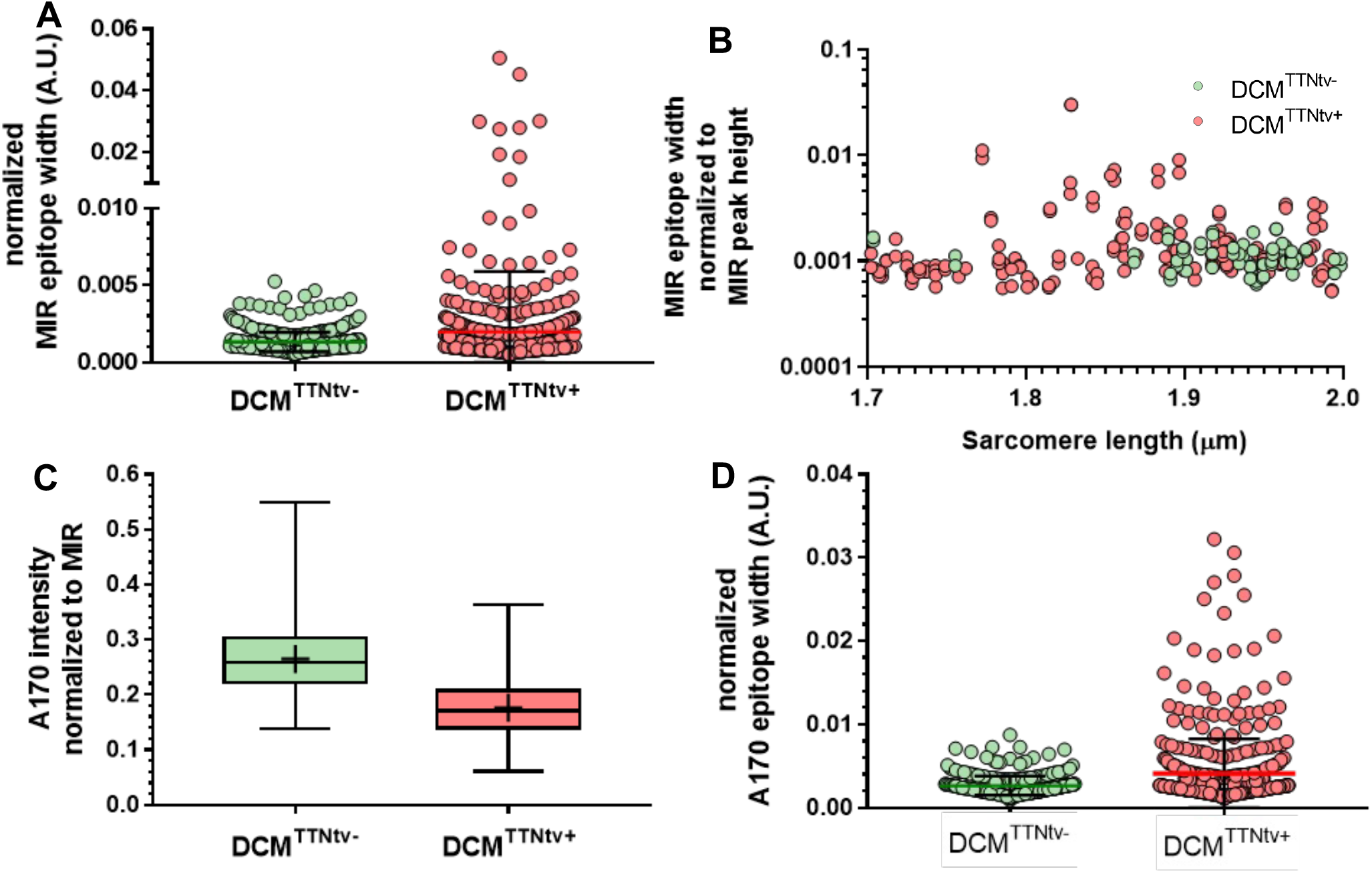
Titin epitope width and intensity analysis. **A,** Full width at half maximum (FWHM) values normalized to peak height of the MIR epitope of DCM^TTNtv−^ and DCM^TTNtv+^ sarcomeres between 1.8 and 2.6 μm. Note that the average sarcomere length values do not differ significantly between the two groups, hence sampling across the investigated sarcomere length range is evenly distributed. Black whiskers: standard deviation; colored, horizontal thick lines: arithmetic mean. **B,** Normalized width distribution of the MIR epitopes in short sarcomeres with lengths between 1.7 and 2.0 μm. **C,** A170 intensity normalized to the peak height of the neighboring MIR epitope within the half sarcomere. Cross marks show the arithmetic mean, the line across the box represents the median. Whiskers indicate the minimum and maximum values, respectively. **D,** FWHM values normalized to peak height of the A170 epitope of DCM^TTNtv−^ and DCM^TTNtv+^ sarcomeres with lengths between 1.8 and 2.6 μm. Note that only the full-length titin of the DCM^TTNtv+^ sarcomeres contain the A170 epitope. Black whiskers: standard deviation; colored, horizontal thick lines: arithmetic mean.

To investigate the structural integrity of the sarcomere-incorporated truncated titin, we carried out measurements on myocardial samples exposed to mechanical stretch. Epitope-to-epitope distance measurements revealed that the A-band-titin length, measured as the distance between two consecutive MIR epitopes separated by an A170 epitope doublet (**Figure 6A**), increased in both groups as the fibers were passively stretched (**Figure 6C**). Regression analysis of the A-band-titin length in the 1.8-2.6 μm sarcomere-length range revealed a significantly decreased slope in DCM^TTNtv+^ samples compared to that in DCM^TTNtv−^ (**Figure 6D**). The MIR epitope was shifted towards the Z-disks at slack sarcomere lengths in DCM^TTNtv+^, and it was less responsive to longitudinal stretch as indicated by the significantly lower A-band-titin length values measured at longer sarcomere values. The average MIR epitope width was significantly increased in the DCM^TTNtv+^ sarcomeres (**Figure 7A**). Furthermore, the spread of the MIR width data points increased upon increasing the sarcomere length from 1.7 to 1.85 μm, then it declined upon further longitudinal sarcomere stretch (**Figure 7B**).

Measuring the distance between consecutive A170 epitopes allowed us to investigate the structural response of the titin kinase (TK) region to mechanical stretch (**Figures 6E and F**). The TK region localizes in the bare zone of the A-band approximately 70 nm from the M-line in both DCM^TTNtv−^ and DCM^TTNtv+^ sarcomeres, calculated as the half value of the distance of two consecutive A170 epitopes. In DCM^TTNtv−^ the M-line to TK distance increased by ~20 nm upon increasing the sarcomere length from 1.8 to 2.6 μm (**Figure 6F**). By contrast, and to our surprise, the TK moved closer to the M-line in DCM^TTNtv+^ sarcomeres when the fibers were passively stretched, indicated by the negative slope of the M-line to TK *versus* sarcomere-length function (**Figure 6F**). The A170 epitope width was significantly greater in the DCM^TTNtv+^ sarcomeres (**Figure 7D**).

## Discussion

Heterozygous titin truncating variants (TTNtv) are the most common genetic cause of familial dilated cardiomyopathy (DCM), accounting for 15-25% of the cases^6, 9, 10^. The pathomechanisms by which titin mutations induce the cardiac phenotype are under extensive research^36^. Although haploinsufficiency and dominant negative effect have recently been suggested based on proteomic analyses^22, 23^, the mechanistic links from the truncated titin protein to sarcomeric structure and function remain highly controversial and debated^37^. In order to dissect titin’s role in the pathogenesis of DCM, here we performed next-generation sequencing, high-resolution protein analysis and super-resolved immunofluorescence microscopy combined with sarcomere extension on cardiac explant samples from a cohort of 127 clinically diagnosed DCM patients.

We identified TTNtvs in 15% of our patient cohort, which is in accordance with prior NGS data^6, 9, 22^. We uncovered additional, non-titin-related DCM and non-DCM causing mutations in the samples (see **Supplementary Tables 2-3**). Clinical data revealed sex differences, as more men carried the truncating mutations than women did. However, based on the echocardiographic measurements, we did not find differences between TTNtv^+^ *versus* TTNtv^-^ DCM patients (**Table 1**). It is important to note that the echocardiographic data were collected just prior to heart transplantation, by which time all of the patients had developed end-stage heart failure. Furthermore, there was a variation in the sample size of the echocardiographic data due to the heterogeneity in clinical profiling. Altogether, there were no significant functional differences between the TTNtv^+^ and TTNtv^-^ DCM patients, which is in line with the recent study of McAfee *et al*^23^.

The evaluation of titin expression revealed increased titin N2BA/N2B ratios in all of the DCM samples compared to healthy donor heart data from literature^34^. We note here that we did not have any non-implanted donor hearts for comparison. However, no differences were seen between the N2BA/N2B ratios in the two DCM groups, suggesting that the more compliant N2BA titin compensates for functional impairment in DCM despite the etiology of the disease^34^. Similarly to the findings of Fomin *et al*^22^ and McAfee *et al*^23^, T1/MyHC was significantly decreased in the DCM^TTNtv+^ group, supporting the tentative hypothesis that haploinsufficiency indeed contributes to the pathomechanism of TTNtv-induced DCM. In addition, we found significantly increased amounts of T2 in the TTNtv+ samples, which points at an increased titin turnover related to the ubiquitin proteasome system, the pathogenic role of which has been suggested by Fomin *et al*^22^, and which needs to be clarified by further experiments. Importantly, however, we found that the integral titin amount, which includes the full-length, truncated and proteolysed proteins, was comparable in the DCM^TTNtv−^ and DCM^TTNtv+^ samples (**Figure 2**). We discovered truncated proteins in the majority (12 out of the 19) of the TTNtv^+^ samples by gel electrophoresis. The truncated proteins were revealed on the gels at the molecular weight levels expected based on the NGS data (**Figures 2A and S1**, **Supplementary Table 4**). The difference in titin expression in DCM^TTNtv−^ and DCM^TTNtv+^ samples, calculated as the T1/MyHC ratio (**Figure 2C**), was alleviated if the truncated proteins were taken into account in calculating total titin in the DCM^TTNtv+^ samples (**Figures 2E and G**). Because the total amount of titin is not significantly reduced in DCM^TTNtv+^, it is unlikely that titin haploinsufficiency plays a decisive role in the pathogenesis of the disease. Rather, the structural and mechanical consequences of the presence of the truncated titin protein must be investigated in detail.

By using Western-blot analysis, we were able to establish that the additional protein bands, identified putatively as the protein products of the truncated titin gene, are indeed truncated titins rather than further degradation products of T2 (**Figure 3**). While the M8M10 antibody, which targets titin near its C terminus, labeled T1 and T2 exclusively (**Figure 3B**), the T12 antibody, which targets titin near its N-terminus, labeled T1 and all the additional protein fragments as well (**Figure 3A**). Thus, the additional protein bands indeed correspond to the protein products of the truncated titin genes. We note here that we could not detect truncated titins in all of the DCM^TTNtv+^ samples (7 out of 19). Conceivably, truncated protein is not produced in all cases^10^. Moreover, the quantity of the truncated proteins was uneven, most likely reflecting an unequal penetrance of TTNtv.

To explore whether the truncated titin is incorporated into the sarcomere, we first analyzed the protein composition of washed myofibrils that are devoid of the sarcoplasm. Electrophoretic analysis of skinned DCM^TTNtv+^ myofibril samples revealed the presence of the respective truncated titin (**Figures 4 and S2**). The results support the findings of Fomin *et al* and McAfee *et al* and suggest a poison peptide mechanism^22, 23^. Fomin *et al* coined that the truncated proteins are accumulated as intracellular aggregates^22^. The study of McAfee *et al* revealed TTNtv variants in sarcomere-containing cellular fractions, suggesting that the truncated titin is incorporated into the sarcomere^23^. However, they could not rule out that the truncated titins are solely present as non-sarcomeric aggregates^23^; therefore, whether the truncated titin molecule is structurally and mechanically integrated into the sarcomere remained a puzzling question.

To uncover the arrangement of truncated titin in the slack and extended sarcomere, we carried out STED super-resolution microscopy on DCM^TTNtv−^ and DCM^TTNtv+^ myocardial tissue samples exposed to mechanical stretch and labeled with sequence-specific anti-titin antibodies (**Figures 5–7**). It is important to note that because the truncated titin molecules do not carry epitopes which are unique with respect to the full-length molecule, the immunofluorescence microscopic results provide only indirect evidence of the truncated titin’s sarcomeric behavior. Since both the full-length and truncated titins are likely present in the sarcomere due to the heterozygous nature of TTNtv, truncated-titin behavior may be inferred from the number, location, intensity and spatial width of the antibody epitope label signals within the sarcomere. We used two anti-titin antibodies to monitor different regions of titin. MIR labels the I/A junction of titin and A170 is localized at the titin kinase (TK) region, at the edge of the bare zone of the A-band. Considering that TTNtv is over-represented in the A-band region, MIR and A170 labeled all and none of the investigated DCM^TTNtv+^ samples, respectively (**Figure 1**). Such a differentiated labeling allowed us to gain precise insight into the sarcomeric behavior of the truncated titin molecules.

We observed no gross structural disturbance in the DCM^TTNtv+^ sarcomeres (**Figure 5**). In fact, based on the overall across-the-sarcomere appearance of both the MIR and A170 epitopes, the myofilaments are in precise registry. We successfully resolved the A170 epitope doublet, with an average separation distance of ~140 nm, owing to the high resolution of STED microscopy which was tested to be ~40 nm in our instrument. Being able to resolve the A170 epitope doublet was important so as to alleviate confounding in intensity measurements and to uncover the behavior of the titin kinase region. The average intensity of the A170 epitope was reduced by ~30% in the TTNtv+ samples while that of the MIR epitope remained unchanged, indicating that the truncated titin is indeed incorporated into the sarcomere, and supporting our results on the protein analysis of washed myofibrils. Furthermore, the lack of MIR epitope doubling suggests that the truncated titin is not only incorporated in the sarcomere but structurally integrated similarly to the full-length form.

The precise epitope localization made possible by STED microscopy allowed us to investigate the structural rearrangements of titin in DCM^TTNtv−^ and DCM^TTNtv+^ sarcomeres exposed to mechanical stretch (**Figures 6–7**). The A-band titin length, measured as the MIR-MIR distance (**Figure 6A**) increased in both groups upon passive stretch, which supports earlier notions that the A-band section of titin is genuinely extensible^38^. Interestingly, however, regression analysis revealed a significantly reduced slope in the TTNtv+ samples across the 1.8-2.6 μm sarcomere range, which points at a reduced A-band extensibility in the DCM^TTNtv+^ sarcomere. Notably, the MIR epitope is shifted towards the Z-disk in the DCM^TTNtv+^ sarcomere at slack (**Figure 6D**), suggesting that A-band titin is more extended, or *vice versa* the I-band titin is more contracted on average (**Figure S3D**). The differences in sarcomere-length-dependent MIR-MIR epitope distance behavior are coupled with a significantly increased MIR epitope width in the DCM^TTNtv+^ sarcomeres (**Figure 7A**), indicating that there is a slight disarrangement among the titin molecules in spite of the gross alignment (i.e., there is no MIR epitope doubling). Presumably, it is the I-band section of the truncated titin molecules that becomes more contracted, owing to the *a priori* weaker A-band attachment, which results in the widening of the MIR epitope. Notably, the average MIR epitope width in the DCM^TTNtv+^ sarcomeres decreases with increasing sarcomere length (**Figure 7B**), suggesting that the axial titin disarrangement may be reduced by mechanical stretch.

The response of the titin kinase region to sarcomere stretch was quite different in the DCM^TTNtv−^ *versus* DCM^TTNtv+^ sarcomeres (**Figure 6F**). While the M-line to TK distance increased with sarcomere length in DCM^TTNtv−^, it progressively decreased in DCM^TTNtv+^. The increase in the M-line to TK distance in DCM^TTNtv−^ provides direct evidence that the TK indeed responds, probably by *in situ* partial unfolding^39^, to mechanical stretch. Notably, the extensibility of the TK region is more than twice as large as in the entire A-band section of titin (**Supplementary Information**), which points at a differential control of titin elasticity or conformation along the thick filament. The reduction in M-line to TK distance in DCM^TTNtv+^ is puzzling, considering that an apparent contraction in the TK region upon sarcomere stretch is unexpected. The controversial TK-region behavior is coupled with a significantly increased A170 epitope width (**Figure 7D**), indicating that there is structural disarrangement among the fewer full-length titin molecules in the bare zone of the DCM^TTNtv+^ sarcomere.

We propose the following model to explain our complex and somewhat puzzling observations (**Figure 8**). In DCM^TTNtv+^, three full-length and three truncated titin molecules per half thick filament, on average, are integrated in the sarcomere. Because the anchorage of TTNtv in the A-band is compromised, the molecules are pulled slightly towards the Z-line by their intact I-band sections. Therefore, in the slack DCM^TTNtv+^ sarcomere the A-band titin length is increased and *vice versa* the I-band titin length is reduced. However, this disposition (and hence the titin disarrangement) is slight, due probably to a pre-stretched state of the A-band section of the truncated titin that enhances its binding within the A-band. The MIR epitopes are slightly out of register, resulting in an increase in the STED epitope profile width. Because TTNtv lacks a good portion of its A-band section and its entire M-band section, only about half of the titins contribute to the A170 signal, hence the A170 epitope intensity is reduced. The reduced number of titins in the bare zone and M-band likely results in structural disarrangement and weakening, leading to an increase in both the M-line to A170 distance (by ~10 nm) and the width of the A170 epitope (**Figure 8B**). This pathological disarrangement is indicated in the figure, albeit in an exaggerated way, by a crooked M-band. Upon stretch (**Figure 8D**), the apparent A-band titin length is increased, but to a smaller degree than in the DCM^TTNtv−^ sarcomere due to the pre-stretched and stabilized nature of the truncated titin molecules. Accordingly, the MIR epitopes on the normal and truncated titin molecules approach each other resulting in a relative narrowing of the STED intensity profile. The M-line to A170 epitope distance becomes reduced upon sarcomere stretch, which is a paradoxical phenomenon due, conceivably, to a mechanically-driven ordering in the M-band. In principle, the faulty mechanosensor function of the M-band, revealed here, may be a major pathway leading to manifest DCM. Although some elements of our proposition, such as the pre-stretched and stabilized A-band portion of the truncated titin and the structurally disarranged M-band, are hypothetical and need further exploration, the model is consistent with our experimental data and provides testable predictions.

**Figure 8.**
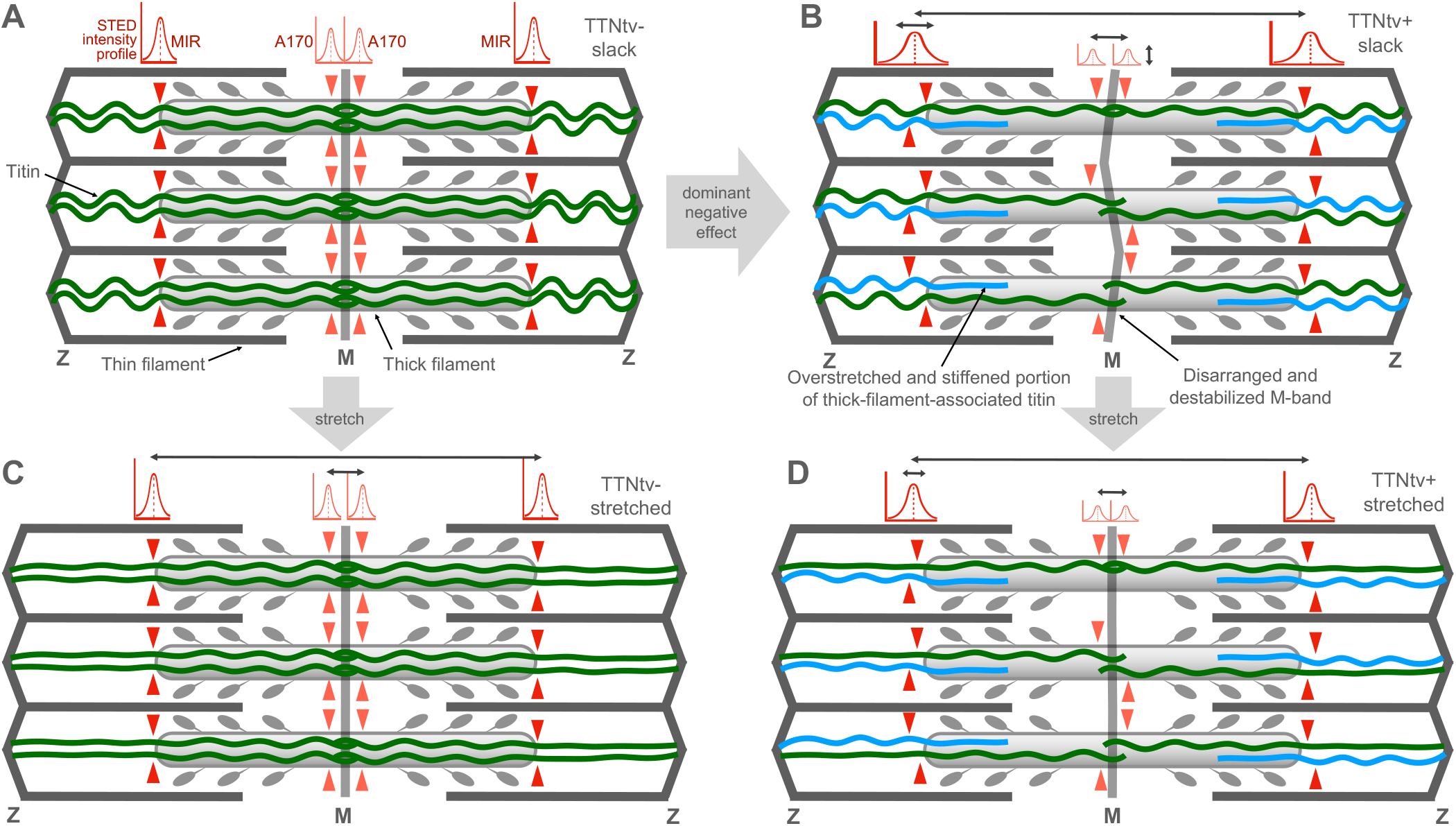
Schematic model of the structural and mechanical changes in the cardiac sarcomere caused by the presence of truncated titin. Gaussian functions above the sarcomere schemes indicate the intensity profiles of the anti-titin epitopes used in this work. The red arrowheads indicate the epitope positions in the respective titin molecules. The black double arrows indicate the changes in measured parameters. **A**, TTNtv-sarcomere at slack. Full-length titin molecules (six per half thick filament, shown as green lines) are present. The anti-titin epitopes are in register and the corresponding STED profiles are of high intensity and have narrow width. **B**, TTNtv+ sarcomere at slack. Truncated titin (shown as blue line) is incorporated in the sarcomere (three per half thick filament on average, presumably randomly distributed). Because the anchorage of TTNtv in the A-band is compromised, the molecules are pulled slightly towards the Z-line (hence the A-band titin length is increased). The MIR epitopes are slightly out of register, resulting in an increase in the STED epitope profile width. Because only about half of the titins contribute to the A170 labeling, the A170 epitope intensity is reduced. The M-line to A170 distance is slightly increased (by ~10 nm), suggesting that the M-band region is disarranged (indicated by a crooked M-band). **C**, TTNtv-sarcomere after stretch. The apparent A-band titin length is increased (by ~236 nm per every μm of sarcomere length increment), but the MIR epitopes remain in register. The M-band to A170 epitope distance is increased upon sarcomere stretch, which is most likely due to structural changes in the titin kinase domain related to its mechanosensory function.^39^ **D**, TTNtv+ sarcomere after stretch. The apparent A-band titin length is increased, but to a smaller degree than in the TTNtv-sarcomere. The MIR epitopes on the normal and truncated titin molecules approach each other resulting in a relative narrowing of the STED intensity profile (see **Figure 6B**). The M-line to A170 epitope distance becomes reduced upon sarcomere stretch due to a mechanically-driven ordering indicated by a straightened M-band.

In conclusion, our results provide strong support to the notion that titin truncating variants are a major cause of familial DCM. However, it is unlikely that titin haploinsufficiency contributes significantly to the pathogenesis of TTNtv-induced DCM. Rather, truncated titin molecules are incorporated and integrated in the sarcomere and likely cause small but significant internal structural and mechanical perturbations. The compensatory effects in the I/A junction and the faulty mechanosensor function in the M-band region of titin are likely to play a significant role in the pathway towards dilated cardiomyopathy.

## Acknowledgements

We gratefully acknowledge the assistance of Krisztina Lór with experimental preparations. She consented to the acknowledgement. We thank Prof. András Csillag and Dr. Gergely Zachar at Semmelweis University, Department of Anatomy, Histology and Embryology for providing access to the Microm HM560 Cryostat.

